# A Combination of Four Nuclear Targeted Effectors Protects *Toxoplasma* Against Interferon Gamma Driven Human Host Cell Death During Acute Infection

**DOI:** 10.1101/2023.12.24.573224

**Authors:** Brittany Henry, L. David Sibley, Alex Rosenberg

## Abstract

In both mice and humans, Type II interferon-gamma (IFNγ) is crucial for regulation of *Toxoplasma gondii* (*T. gondii*) infection, during acute or chronic phases. To thwart this defense, *T. gondii* secretes protein effectors hindering the host’s immune response. For example, *T. gondii* relies on the MYR translocon complex to deploy soluble dense granule effectors (GRAs) into the host cell cytosol or nucleus. Recent genome-wide loss-of-function screens in IFNγ-primed primary human fibroblasts identified MYR translocon components as crucial for parasite resistance against IFNγ driven vacuole clearance. However, these screens did not pinpoint specific MYR-dependent GRA proteins responsible for IFNγ signaling blockade, suggesting potential functional redundancy.

Our study reveals that *T. gondii* depends on the MYR translocon complex to prevent host cell death and parasite premature egress in human cells stimulated with IFNγ post-infection, a unique phenotype observed in various human cell lines but not in murine cells. Intriguingly, inhibiting parasite egress did not prevent host cell death, indicating this mechanism is distinct from those described previously. Genome-wide loss-of-function screens uncovered TgIST, GRA16, GRA24, and GRA28 as effectors necessary for a complete block of IFNγ response. GRA24 and GRA28 directly influenced IFNγ driven transcription, GRA24’s action depended on its interaction with p38 MAPK, while GRA28 disrupted histone acetyltransferase activity of CBP/p300. Given the intricate nature of the immune response to *T. gondii*, it appears that the parasite has evolved equally elaborate mechanisms to subvert IFNγ signaling, extending beyond direct interference with the JAK/STAT1 pathway, to encompass other signaling pathways as well.

## Introduction

*Toxoplasma gondii* (*T. gondii*), an apicomplexan intracellular parasite, is prevalent among warm-blooded animals, impacting around one-third of the worldwide human population (1). The majority of isolates in North America and Europe belong to one of three distinct lineages where type I strains are acutely virulent, while type II are intermediate and type III are avirulent in the laboratory mouse (2). While most infections are asymptomatic, toxoplasmosis can pose significant risks for immunocompromised patients and infants infected in utero (3). The reactivation of chronic infections plays a pivotal role in the progression of the disease, as the parasite transitions from the lytic tachyzoite stage, characteristic of acute infections, to the slowly growing bradyzoite stage (4). Upon invading the host cell, the parasite establishes a replication-permissive niche within the parasitophorous vacuole (PV), a compartment isolated from the host cytoplasm by the parasitophorous vacuole membrane (PVM) (5).

*T. gondii* employs a multifaceted strategy to manipulate various host cell functions, achieved through the secretion of numerous effectors originating from rhoptries and dense granules into the host cell (6). Dense granule proteins (GRAs) are secreted once the parasites have successfully entered the vacuole and may be directed to the PVM (i.e., GRA15, MAF1), the host cell cytoplasm (i.e., GRA18, MAG1), or the host cell nucleus (i.e., GRA16, GRA24, GRA28, HCE1/TEEGR, TgIST and TgNSM) (7). The successful traversal of the PVM by GRA effectors is facilitated by a translocon complex located on the PVM and comprised of MYR proteins: MYR1, MYR2, MYR3, MYR4, and ROP17 (8–11), as well as GRA45, and ASP5 that are responsible for GRA protein processing and trafficking to the PVM. (12, 13).

In both mice and humans, Type II interferon-γ (IFNγ) is essential for immune control of *T. gondii* infection, in both acute and chronic phases of infection (14, 15). Decreased IFNγ levels have been observed in AIDS patients with cerebral toxoplasmosis (16, 17), highlighting the significance of this immune response. IFNγ triggers JAK1 and JAK2 activation, resulting in the phosphorylation of STAT1 homodimers and the initiation of gene expression from core promoters containing a gamma-activated sequence (GAS) (18).

*T. gondii* disrupts host interferon responses by manipulating epigenetic gene regulation through two secreted effectors, *Toxoplasma* inhibitor of STAT1 transcriptional activity (TgIST) (19, 20) and *Toxoplasma* NCoR/SMRT modulator (TgNSM) (21), which impact host corepressor complexes Mi-2/NuRD and NCoR/SMRT respectively. TgIST sequesters STAT1 and recruits NuRD, leading to the formation of nonpermissive chromatin along with prevention of recruitment of the histone acetyltransferase coactivators CREB-binding protein (CBP) and EP300 (p300), to STAT1 thereby preventing subsequent transcriptional activation of ISGs (22). TgNSM elevates NCoR/SMRT levels, enhancing its repressive activity and blocking a subset of interferon dependent genes (21). TgIST is crucial for *T. gondii*’s virulence at all infection stages, while TgNSM acts primarily to protect intracellular cysts from premature host cell death and rupture. Together, they thwart the expression of interferon-regulated necroptotic genes, preventing host cell death infected with the latent-stages bradyzoites (23).

In mice, the antiparasitic effects of IFNγ are mediated by the activation of a subset of interferon-stimulated genes (ISGs), which include immunity-related GTPases (IRGs) and guanylate binding proteins (GBPs) (24, 25). These proteins are recruited to the PVM, leading to its disruption and parasite death (26–28). To evade this host defense, *T. gondii* deploys the ROP5/ROP18/ROP17 complex to deactivate IRG proteins, thus preventing PVM damage (Hunter and Sibley, 2012; Mukhopadhyay et al., 2020).

Humans lack functional IFNγ inducible IRGs, and the mechanisms involved in IFNγ-mediated restriction of *T. gondii* in humans tend to vary depending on the specific cell type and the strain of *Toxoplasma* involved. These mechanisms include processes such as nutrient deprivation (29), endo-lysosomal destruction (30, 31), autophagy (32), and induction of host cell death (33). A comprehensive gain of function screen of hundreds of ISGs induced by IFNγ and a genome wide loss of function screen conducted in human cells uncovered only a few host effectors capable of restricting *T. gondii* growth including RARRES3 that acts only against type III strains (34) and the E-3 ubiquitin ligase RNF213 that acts against multiple strain types in a range of cell types (35). These findings emphasize the complexity of immunity to *T. gondii*, indicating that a multitude of factors are likely involved in influencing the outcome of infection.

In accord with this conclusion recent genome wide loss of function screens in *T. gondii* have identified multiple genes critical for the parasite fitness in IFNγ stimulated human foreskin primary fibroblasts (HFFs) (36–38). A comparison of gene sets associated with *T. gondii* fitness in screens conducted in-vivo (39–41) or in IFNγ-stimulated murine cells (13) with those from human cells reveals a substantial discrepancy between the two hosts, suggesting variations in parasiticidal mechanisms. Remarkably, in these screens, only the disruption of MYR traslocon complex genes (MYR3, MYR1, and ROP17), which resulted in the complete cessation of all GRA secretion beyond the PVM, showed a survival phenotype. In contrast, none of the individual GRA genes, including those with recognized anti-IFNγ properties (e.g., TgIST, TgNSM), displayed significant phenotypes in those screens. These results suggest that *T. gondii* may depend on the combined function of multiple nuclear-targeted effectors to efficiently block the IFNγ response, and the importance of these effectors could vary between murine and human hosts.

In our current study, we uncover that *T. gondii* depends on the MYR translocon complex to prevent host cell death and premature egress in human cells stimulated with IFNγ post infection. By employing a genome-wide loss-of-function screens, we have deconvolved the roles of four separate effectors-TgIST, GRA16, GRA24, and GRA28 as the critical MYR dependent effectors responsible for blocking the IFNγ response. This reveals that the parasite’s strategies to subvert IFNγ signaling go beyond direct interference with the JAK/STAT1 pathway, extending to the manipulation of additional biological pathways.

## Results

### MYR1 prevents tachyzoite early egress and host cell death stimulated with IFNγ following infection

During the characterization of host IFNγ response to acute stages of *T. gondii* infection (tachyzoites), we discovered that IFNγ treatment of HFFs post infection with RH type I (RH) or type II (ME49) *Δmyr1* translocon mutants, led to early parasite egress (Fig. 1A, Fig. S1A). In these experiments, we subjected naive HFFs to IFNγ stimulation four hours after infection with either WT, *Δmyr1*, or *Δmyr1*::MYR1 complemented parasites, and subsequently assessed them via immunofluorescence analysis 24 hours post-infection. While the majority of WT parasites appeared to reside within intact vacuoles, vacuoles of Δmyr1 mutants were observed to be ruptured (Fig 1B, Fig. S1A). This early release led to the presence of multiple singly reinvaded parasites, ultimately resulting in a higher number of vacuoles of significantly reduced size. (Fig. 1C, 1D, Fig. S1B, S1C). Premature egress is frequently triggered by the activation of various cell death pathways, each governed by distinct mechanisms (21, 33, 42, 43). If the initiation of host cell death serves as trigger for premature egress, we would expect to observe host cell death even when parasite egress is blocked. Therefore, to assess the effect of MYR1 on host cell survival, we directly measured the kinetics of RH Δmyr1-mCherry egress from HFFs via time-lapse imaging of live infected cells by video microscopy. Four hours after infecting HFFs with *Δmyr1*-mCherry mutants, the cells were left untreated or subjected to stimulation with either IFNγ alone or in combination with Compound 1, a potent inhibitor of Toxoplasma gondii protein kinase G (PKG), essential for parasite egress (44, 45). Upon IFNγ treatment, parasites exhibited significantly earlier egress compared to the control group (Fig. 1E,F. Movie S1, S2). However, when Compound 1 was introduced during infection, the majority of infected cells displayed characteristics such as cell rounding, detachment, and the uptake of live-dead stain SYTOX green (Fig. 1E,F, Movie S3, S4). By the 24-hour mark following IFNγ treatment, 100% of the parasites had already undergone egress. In contrast, host cells treated with IFNγ and Compound 1 displayed early indicators of cell death at the same time point, ultimately resulting in a significant 60% mortality rate among infected host cells after 60 hr of continuous time-lapse observation. (Fig. 1E,G, Movie S3, S4). Next, we asked whether chemical inhibitors of cell death pathways could reduce IFN-γ-driven premature egress. Among these inhibitors were the pan-caspase inhibitor Z-VAD-FMK, which blocks apoptosis; the necroptosis inhibitors GSK’963, GSK’872, and necrosulfonamide (NSA), which block RIPK1, RIPK3, and MLKL activity, respectively; and the pyroptosis inhibitor Z-YVAD-FMK, which blocks caspase 1. We also tested the effect of autophagy inhibitor 3-methyladenine (3-MA) which blocks PI3K and ferroptosis inhibitor Ferrostatin-1 (Fer-1). To assess the impact of these inhibitors on the early egress phenotype and host cell viability, we utilized the quantification of lactate dehydrogenase (LDH) release into the culture supernatant. The addition of these inhibitors, either individually or in combination, concomitant with IFNγ, did not rescue the early egress phenotype (Fig. S1D). Given IFNγ known role in inducing tryptophan degradation in human fibroblasts (Pfefferkorn, 1984) and the association of tryptophan deficiency with cell death (Taylor et al., 1996), we assessed whether the addition of tryptophan could rescue the early egress phenotype. Supplementing the media with tryptophan upon IFNγ stimulation did not block cell death as well (Fig. S1D). As cell death pathways often intersect, it remains plausible that chemical inhibition may be incapable of halting previously initiated cell death from diverting along an alternative pathway triggering parasite egress. Alternatively, multiple cell death pathways could be activated simultaneously.

**Figure 1.**
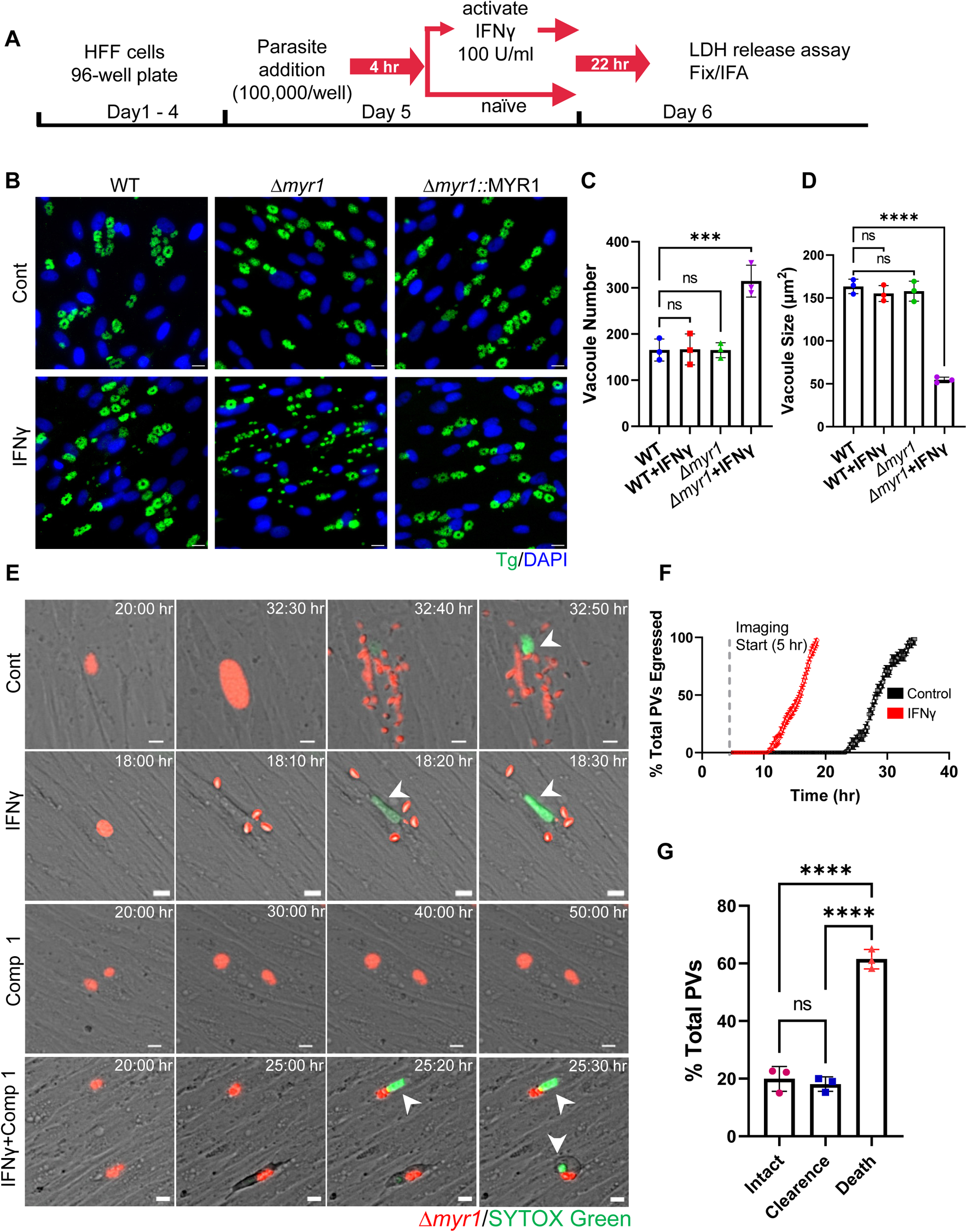
IFNγ stimulation following infection is countered by MYR1, preventing early tachyzoite egress and host cell death. (A)Schematic illustration of the workflow used to examine tachyzoite growth dynamics. HFF cells were inoculated into 96-well plates and allowed to reach confluency prior to the addition of parasites (100,000/well). Parasites were allowed to invade for 4 hr prior to ± IFNγ 100 U/ml treatment for 22 hr followed by LDH release assay, fixation and IF staining. (B) Representative images of HFFs infected with RH (WT), *Δmyr1* and *Δmyr1*::MYR1 complement mutants and treated as in (A). Cells were labelled with DAPI for nuclei (blue) and anti-GAP45 for parasites (green). Scale bar = 20 µm. (C) Average PV number per field. (D) Average PV size. Data in (C) and (D) represent means ± SD of three biological replicates conducted in technical duplicate with at least 30 images per sample and replicate. Statistical significance was determined using one-way ANOVA with Dunnett’s multiple comparison test. ***P<0.001, ****P<0.0001. (E) Time-lapse images of HFFs infected for 4 hr with RH *Δmyr1*-mCherry mutants (red) prior to ± IFNγ 100 U/ml treatment in the presence of STOX green (150 nM) (green) combined with ± 5 µM Compound 1. Live infection was imaged every 10 min starting 5 hr postinfection until 60 hr postinfection. Scale bar = 5 µm. (F) Time of ± IFNγ stimulated *Δmyr1*-mCherry parasites egress was recorded for at least 100 PVs per condition per replicate. The percentage of total parasites egressed by the end of each hour is indicated. (G) Quantification *Δmyr1*-mCherry parasites and infected host cell fate in IFNγ stimulated cells in presence of Compound 1 (5 µM). Data from three independent experiments were pooled. Mean ± SD (n = 3 experiments, 100 parasite vacuoles were counted in each treatment). Statistical significance was determined using one-way ANOVA with Tukey’s multiple comparison test. **** P < 0.0001.

### IFNγ driven early egress phenotype is unique to human cells

To test if the early egress phenotype of *Δmyr1* mutants was exclusive to primary fibroblasts or occurred in other human cell lines, we examined LDH release upon infection with *Δmyr1* mutants followed by IFNγ treatment. The tested cell lines originated from foreskin, lung, intestine, or brain (e.g., HFF-hTERT, A549, HCT-8, and SH-SY5Y), and included monocyte-derived macrophages (THP-1). All but THP-1 cells showed a significant increase in LDH release driven by the early egress phenotype following infection and IFNγ treatment (Fig. 2A).

**Figure 2.**
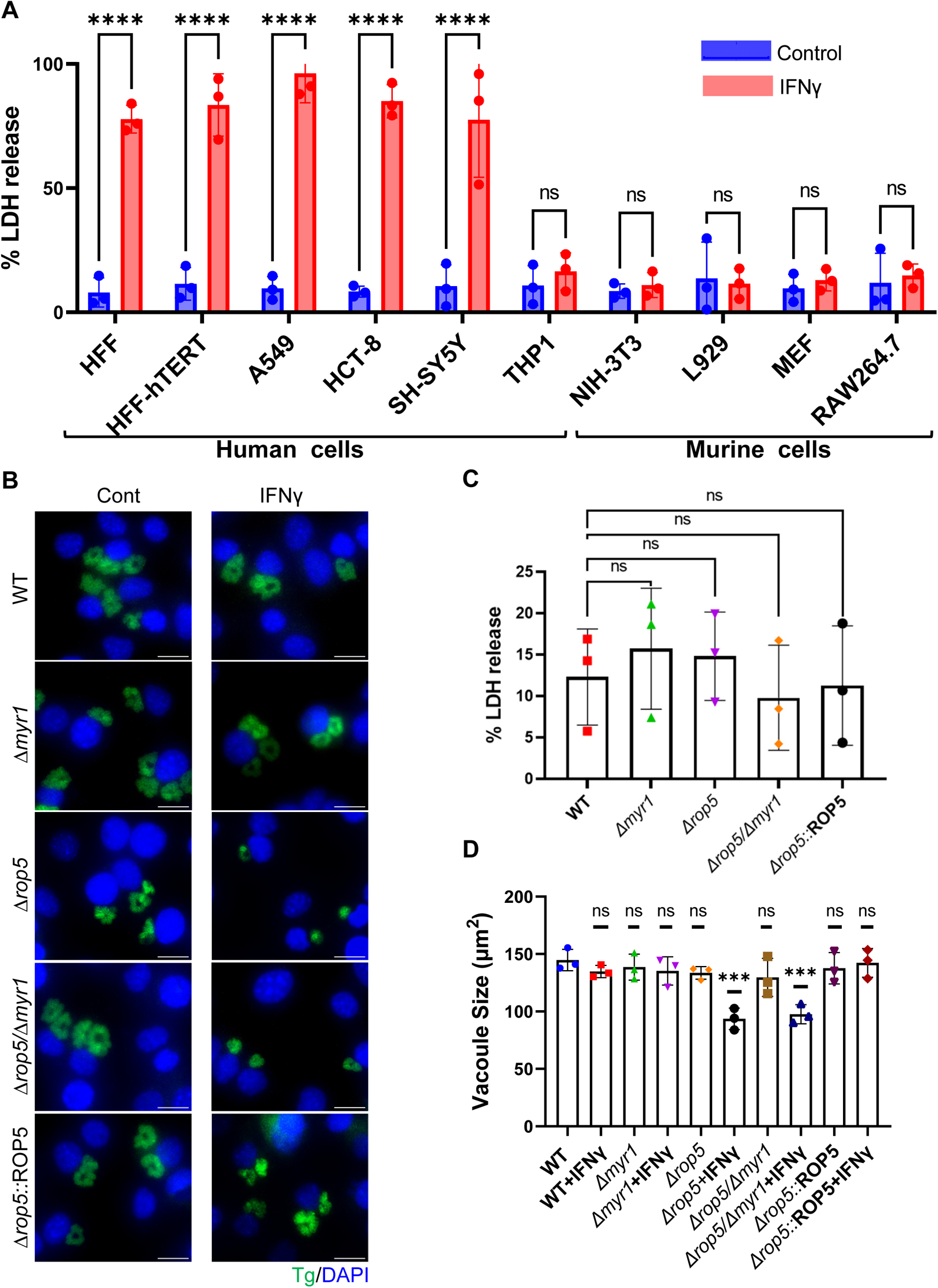
The early egress phenotype induced by IFNγ is distinct to human cells. (A) Human: HFF, HFF-hTERT, A549, HCT-8, SH-SY5Y, THP-1 and murine: NIH-3T3, L929, MEFs, RAW264.7 cell lines were infected with RH *Δmyr1* mutants for 4 hr prior to ± IFNγ 100 U/ml treatment. Cell supernatant was collected 26 hr after infection LDH activity was determined to measure cell lysis. LDH release was calculated as the percent of maximal LDH release (after triton treatment of cells). Data from three independent experiments were pooled. Mean ± SD (n = 3 experiments, each with 3 technical replicates counted in each treatment). **** P < 0.001, one-way ANOVA test with Sidak’s multiple comparison test. (B) Representative images of NIH-3T3 cells infected with RH, *Δmyr1* and *Δmyr1*::MYR1 complement parasites for 4 hr prior to ± IFNγ 100 U/ml treatment. Twenty-six hours post infection cells were fixed and labelled with DAPI for nuclei (blue) and anti-GAP45 for parasites (green). Scale bar = 20 µm. (C) LDH release in the supernatant of NIH-3T3 cells was measured 26 hr post infection. Plotted is the percent of LDH release compared to maximal LDH release (after triton treatment of cells). (D) Average PV size. Data in (C) represents Mean ± SD of three biological replicates Mean ± SD (n = 3 experiments, each with 3 technical replicates counted in each treatment). Statistical significance was determined using one-way ANOVA test with Dunnett’s multiple comparison test. Data in (D) represents Mean ± SD of three biological replicates conducted in technical duplicate with at least 30 images per sample and replicate. Statistical significance was determined using one-way ANOVA with Dunnett’s multiple comparison test. ***P<0.001.

To examine whether an inability to secret GRA proteins beyond the PVM would result in a similar premature egress phenotype in murine cells, *Δmyr1* mutants were used to infect different murine fibroblast cell lines (e.g.NIH-3T3, L929 and MEFs). We also tested the effect of IFNγ stimulation post infection in the murine macrophage cell line, RAW264.7. Surprisingly, we did not detect early egress in any of these cell lines (Fig. 2A).

IFNγ driven mechanisms that kill *Toxoplasma* in mice and murine cells are primarily driven by the activation of Immunity-Related GTPases (IRGs) and Guanylate Binding Proteins (GBPs), which target and destroy the PV and, consequently, the parasite itself. *Toxoplasma* counteracts IRG function by employing the kinase activity of secreted ROP5/ROP18/ROP17 complexes (27, 46). ROP5/ROP18/ROP17 have been shown to be epistatic to the MYR1-dependent TgIST function within the IFNγ pathway (19, 20). Therefore, we tested if genetic depletion of ROP5 locus, the dominant component within the ROP5/ROP18/ROP17 complexes, will unmask the *myr1* dependent phenotype. However, infecting either NIH-3T3 or L929 cells with *Δmyr1/Δrop5* mutants, followed by IFNγ treatment, did not result in early egress, as evidenced by the absence of an increase in LDH release (Fig. 2C, Fig. S2B). Instead, it resulted in a notable reduction in vacuole size (Fig. 2B, Fig. S2A). Interestingly, there was no additional impact on vacuole size when comparing the single ROP5 locus mutant to the double *Δmyr1/Δrop5* knockout. Hence, it appears that *T. gondii* inability to alter host cell signaling upon IFNγ stimulation, achieved through GRA protein secretion beyond the PVM, results in host cell death and premature egress, a phenotype unique to human cells.

### Toxoplasma relies on TgIST, GRA16, GRA24 and GRA28 to block IFNγ driven premature egress during acute infection

We have recently shown that *T. gondii* bradyzoites depend on the combined action of TgIST and TgNSM to block IFNγ signaling and prevent necroptotic host cell death (21). Surprisingly, when assessing LDH release levels post IFNγ stimulation in HFF cells infected with WT, *Δist, Δnsm, Δist/Δnsm*, or *Δmyr1* mutants, we found that only the *Δist* or *Δist/Δnsm* mutants, in addition to Δ*myr1*, resulted in increased LDH release (Fig. 3A). However, this increase amounted to only ∼20% of that detected in *Δmyr1* infected cells (Fig. 3A), suggesting that tachyzoite stages rely on an additional effector, other than TgNSM, that modulates similar pathway but in bradyzoite infected cells.

**Figure 3.**
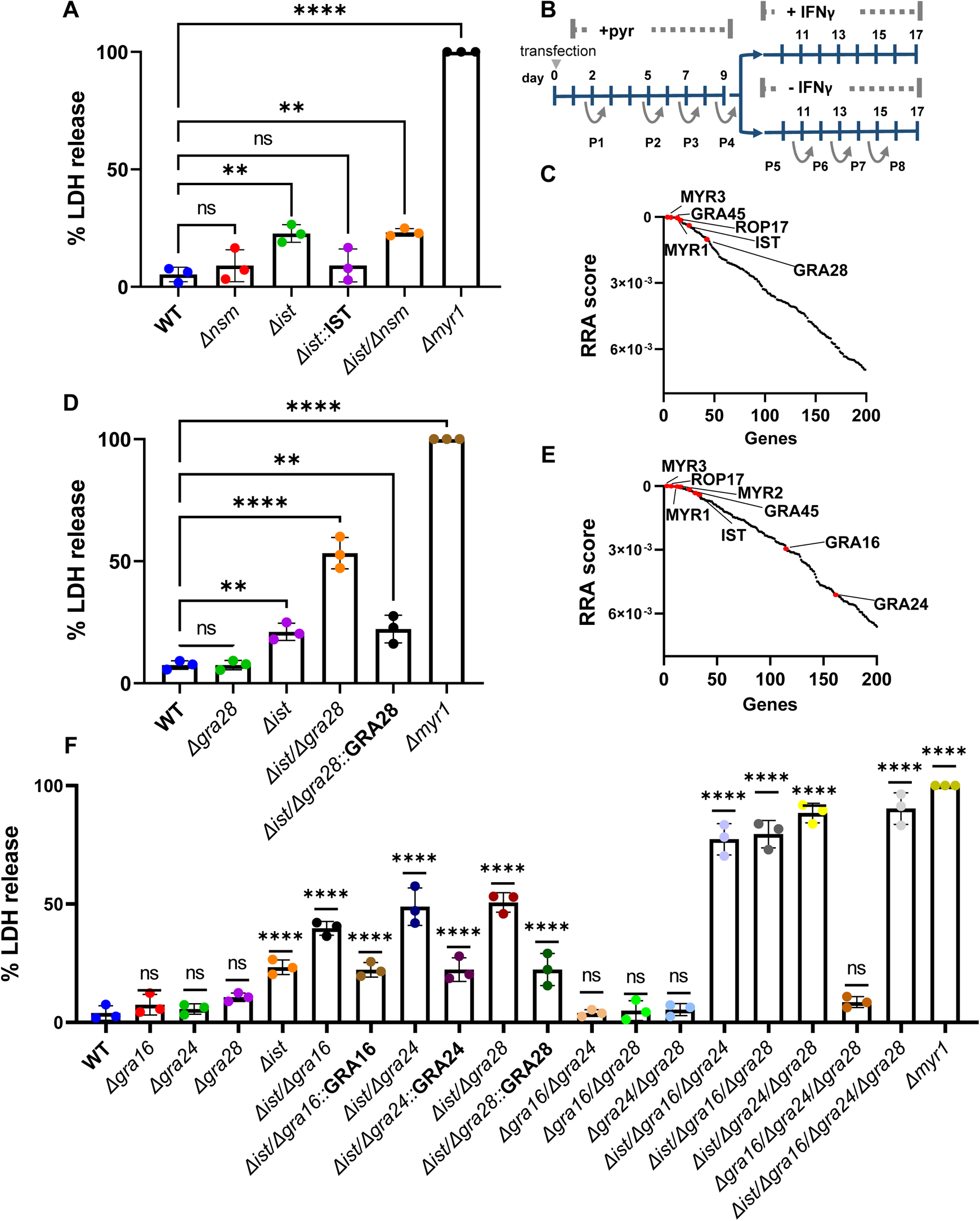
*T. gondii* tachyzoites depend on TgIST, GRA16, GRA24 and GRA28 to block IFNγ driven premature egress. (A) HFFs were infected with RH (WT), *Δnsm*, *Δist*, *Δist*::IST, Δist/Δnsm and *Δmyr1* mutants for 4 hr prior to ± IFNγ 100 U/ml treatment. Cell supernatant was collected 26 hr after infection and LDH activity was determined to measure cell lysis. (B) Screening Process: RH-Cas9 parasites underwent transfection with linearized plasmids housing 10 sgRNAs targeting each *Toxoplasma* gene. The transfected parasites were then propagated through four passages in HFFs under pyrimethamine selection to eliminate non-transfected parasites and those with integrated plasmids containing sgRNAs targeting genes crucial for fitness in HFFs. Following this, the pool of mutant parasites underwent four additional passages, either in naive or IFNγ-stimulated HFFs, 4 hours post-infection. (C) The RRA score distribution of top 200 negative selected genes (IFNγ vs naϊve)), reported by the MAGeCK algorithm. (D) LDH release in HFFs infected with RH (WT), *Δist, Δgra28, Δist/Δgra28, Δist/Δgra28::*GRA28 complement and *Δmyr1* mutants for 4 hr prior to prior to ± IFNγ 100 U/ml treatment was determined to measure cell lysis. (E) RRA score distribution of top 200 negative selected genes (IFNγ vs naϊve) in a screen performed in the RH cas9 *Δgra28* mutant. (F) LDH release in HFFs infected with various RH mutants for 4 hr prior to ± IFNγ 100 U/ml treatment was determined to measure cell lysis. LDH release was calculated as percent of LDH release in *Δmyr1* mutant (A, D, F) and represents Mean ± SD of three biological replicates Mean ± SD (n = 3 experiments, each with 3 technical replicates counted in each treatment. The mean values for the mutants were compared with that for WT using one-way ANOVA with Dunnett’s multiple comparison test; **P < 0.01, ****P<0.0001.

To identify the effector that functions together with TgIST to subvert the IFNγ response we performed a genome-wide loss-of-function screen. To accomplish this, we utilized the RH type I parasite strain engineered to express Cas9 (RH-Cas9) (47). Our approach involved creating a mutant pool of *Toxoplasma* by transfecting RH-Cas9 parasites with a library of sgRNAs, each library containing ten guides designed for each of the 8,156 Toxoplasma genes (48). We allowed this mutant pool to grow for four passages in HFFs. Subsequently, we passed this pool of parasite mutants an additional four times, this time in HFFs that were either infected and left unstimulated or infected and stimulated with IFNγ 4 hr post infection (Fig. 3B). We then conducted the amplification, sequencing, and quantification of the sgRNAs from passage 8 in either naïve or IFNγ-stimulated HFFs. We used the robust rank aggregation (RRA) MAGeCK test (49) to identify genes that were negatively selected under IFNγ treatment. The top negatively selected genes were *MYR3*, *GRA45*, *MYR1* and *ROP17*, all of which affect GRA export (Fig. 3C, Table S1). As expected *TgIST* fell within the top fifteen negatively selected genes. Further examination of the top 200 hits revealed *GRA28* ranking in the thirty-sixth position (Fig. 3C, Table S1). GRA28 is MYR1 dependent secreted effector known to localize within the host nucleus and induce chemotactic migration of parasitized macrophages (50, 51). This discovery prompted us to investigate its potential role in shaping the response to IFNγ. The single deletion of *GRA28* did not result in increased LDH release upon IFNγ treatment (Fig. 3D). However, when both *GRA28* and *TgIST* were deleted, there was a ∼30% increase in LDH release compared to the single *Δist* mutant, amounting to ∼50% of the LDH release observed in the *Δmyr1* infection. This phenotype was rescued by complementing the *Δist/Δgra28* mutant with GRA28 (Fig. 3D).

The partial LDH release phenotype observed in the *Δist/Δgra28* mutant once again suggested the presence of at least one more effector employed by tachyzoites to modulate the host response to IFNγ. To uncover this missing effector, we performed an additional genome-wide loss-of-function screen. For this screen we utilized a RH-Cas9 *Δgra28* strain generated by deleting *GRA28* in the original Cas9 expressing line. By conducting the screen in a *Δgra28* background, we aimed to sensitize the parasites and potentially unveil the elusive missing effectors. In the new screen, the top negatively selected genes were *MYR3*, *GRA45, MYR1*, *ROP17* and *TgIST* (Fig. 3E, Table S2). Further inspection of the top 200 negatively selected genes under IFNγ treatment revealed two other secreted GRA effector genes *GRA16* and *GRA24* (52, 53) (Fig. 3E, Table S2). Single deletion of either *GRA16* or *GRA24* did not result in significantly altered LDH release response to IFNγ post infection compared to the WT (Fig. 3F). Nonetheless, when either *GRA16* or *GRA24* was deleted in the *Δist* background, it resulted in ∼20% and ∼30% increase in LDH release compared to the single *Δist* mutant respectively. Complementing the *Δist/Δgra16* and *Δist/Δgra24* mutants with GRA16 and GRA24 respectively rescued these phenotypes (Fig. 3F). In contrast, other double knockout combinations (i.e., *Δgra16/Δgra24, Δgra16/Δgra28, Δgra24/Δgra28)* exhibited IFNγ responses similar to those of the WT (Fig. 3F). When we assessed triple knockout mutants’ response to IFNγ stimulation (i.e., *Δist/Δgra16/Δgra24, Δist/Δgra16/Δgra28, Δist/Δgra24/Δgra28 and Δgra16/Δgra24/Δgra28*) we observed that all but *Δgra16/Δgra24/Δgra28* mutants had an elevated LDH release reaching ∼70-80% the of that seen in the *Δmyr1* infection (Fig. 3F).The quadruple knockout *Δist/Δgra16/Δgra24/Δgra28* reached ∼90% of LDH release. Collectively, these results indicate that *T. gondii* tachyzoites rely on a combinatorial inhibitory function of TgNSM, GRA28, GRA16 and GRA24 to prevent IFNγ induced premature egress.

### GRA28 and GRA24 affect IFNγ driven transcriptional response

The observed reliance of *T. gondii* on the combined action of TgIST, GRA28, GRA16, and GRA24 to prevent premature egress upon IFNγ stimulation led us to investigate whether these effectors could directly influence the regulated transcription of IFN-γ-responsive genes. To eliminate potential confounding effects from each of these secreted effectors, present during infection, we co-expressed luciferase reporter downstream of GAS sequences in the presence or absence of the four effectors in different combinations in HeLa cells. Consistent with previous reports, the presence of TgIST had an inhibitory effect on the transcriptional activity of the reporter upon IFN treatment (Fig. 4A). GRA28 also exhibited an inhibitory effect, whereas GRA24 enhanced the responsiveness and GRA16 did not significantly affect IFNγ driven transcription. GRA24 modulates cytokine production by interacting with p38α MAPK through its R1 and R2 domains, which contain kinase interaction motifs. (53, 54). Removal of GRA24 R1/R2 domains abolished its ability to induce IFNγ hyperresponsiveness (Fig. 4A). Interestingly, when either TgIST or GRA28 was present with GRA24, it negated the hyperactivation effect of GRA24. Furthermore, the combined expression of all four effectors resulted in an overall inhibitory effect as well (Fig. 4A).

**Figure 4.**
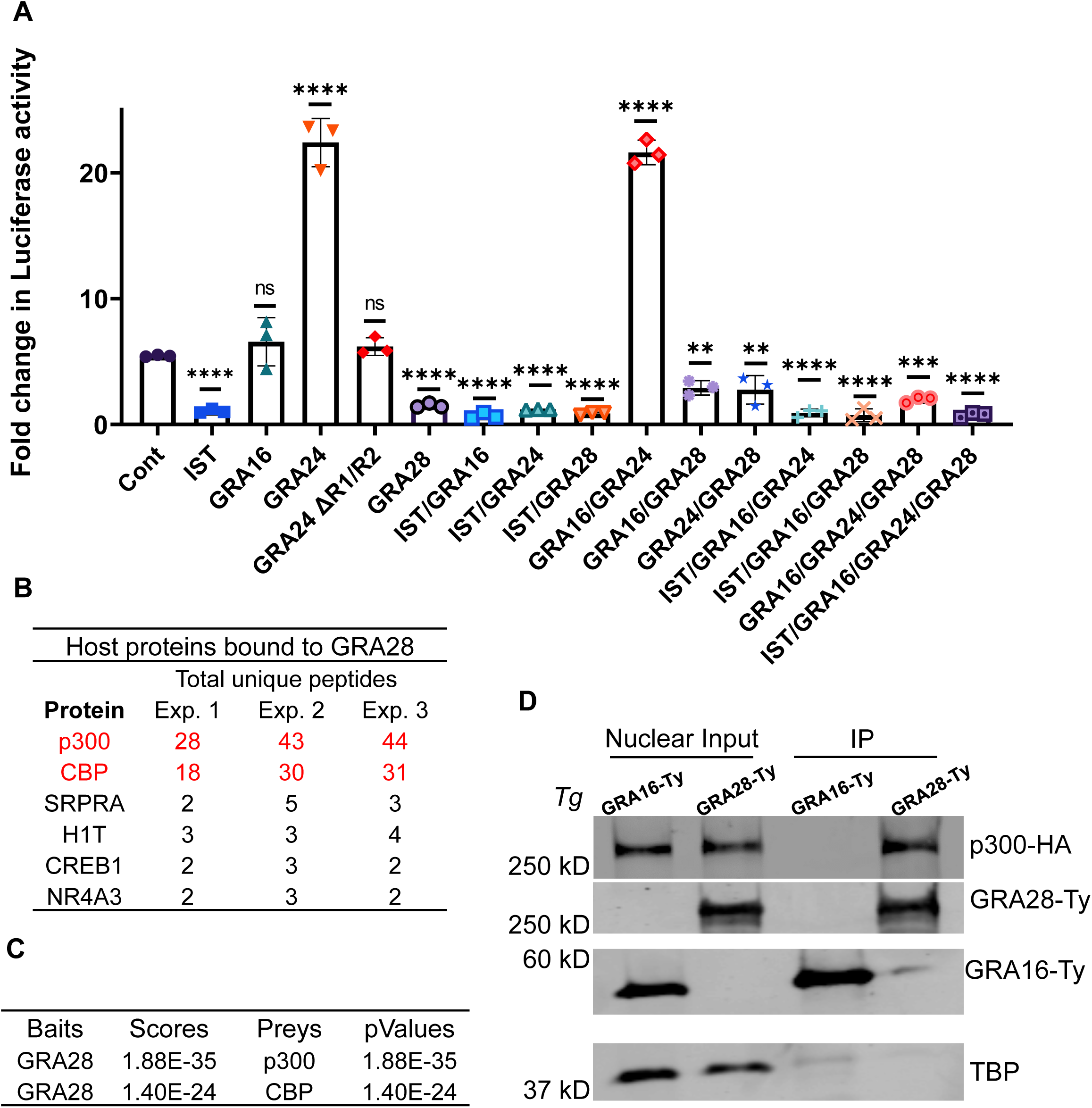
GRA28 targets CBP/p300 histone acetyltransferase and blocks IFNγ driven transcriptional response. (A) GAS luciferase reporter constructs were transiently transfected into HeLa cells with different combinations of IST, GRA16, GRA24, GRA24 ΔR1/R2, GRA28, or empty vector. Twenty-four hours later, transfected cells were treated with IFNγ at (100 U/mL for 24 hr) and firefly luciferase activity was determined. The transfection efficiency was normalized against the-Rennila luciferase activity from the cotransfected pRL-TK vector. Results shown are fold induction over unstimulated control and represent the averages from 3 biological replicates done in duplicate. Each point represents a biological replicate. Mean ± SD (n = 3 experiments, each with 2 replicates). The mean values for the mutants were compared with that for WT using one-way ANOVA with Dunnett’s multiple comparison test; **P < 0.01, ****P<0.0001. (B) GRA28Ty-associated host proteins identified by mass spectrometry (MS) analysis. GRA28-Ty immunoprecipitated from RH GRA28-Ty vs. wildtype untagged RH parasite infected HFF cells. Combination of three experiments showing proteins that were solely identified in GRA28-Ty and not in wildtype infected cells. Identity of the proteins with their respective number of peptides are indicated, CBP and p300 are in red. (C) Three replicates of MS analysis data sets generated with GRA28-Ty and WT IP experiments were analyzed using SFINX to filter out false-positive interactions and rank true-positives. CBP and p300 were identified as GRA28 interactors with high statistical confidence. (C) Western blot analysis of GRA28-Ty and GRA16-Ty immunoprecipitated from nuclear lysates of HEK 293T cells that were transfected with p300-HA. Twenty-four hours after transfection, cells were infected with RH either expressing Ty-tagged GRA28 or GRA16 for 16 hr. The nuclear protein fractions were resolved on SDS-PAGE gels and different proteins were probed with primary antibodies and imaged with LICOR specific secondary antibodies. Equal amounts of nuclear lysates used for immunoprecipitation are loaded alongside as nuclear input controls. Images represent one of two blots performed, all showing similar results.

### GRA28 targets CBP/p300 histone acetyltransferase activity

Considering the results of our CRISPR screen and the transcriptional reporter assay, it became apparent that GRA28 is the second major effector, in addition to TgIST, that *T. gondii* relies on to impede the IFNγ response. Consequently, we directed our focus toward investigating the molecular mechanisms underlying GRA28’s function. Recent studies have revealed that GRA28 associates with host chromatin modifiers including the NuRD and SWI/SNF complexes, resulting in the upregulation of CCR7 expression, which plays a crucial role in the chemotactic migration of infected macrophages (50). In this study, the host protein targets of GRA28 were identified through immunoprecipitation (IP) experiments performed in the murine macrophage cell line RAW264.7. Given that we did not observe a premature egress phenotype in either murine or human macrophages (Fig. 2A), we hypothesized that GRA28 might interact with different host targets depending on the cell type and species. Therefore, we conducted IP experiments using a GRA28-Ty tagged line in HFF cells that were infected with either GRA28-Ty *T. gondii* or WT tag-free parasites for 24 hours. Nuclear extracts were prepared, GRA28-Ty was immunoprecipitated, and complexes from three independent replicates underwent MS/MS analysis. By excluding all host proteins that appeared in the control fraction (Table S3), we were left with a list of only six proteins specifically immunoprecipitated with GRA28-Ty in HFFs. Histone acetyltransferase (HAT) CBP and its homologue p300 exhibited the highest fold increase in unique peptide detection, with 38-fold and 26-fold increases, respectively (Fig. 4B). Further analysis of the MS data sets using the Straightforward Filtering Index program (SFINX) (55) identified p300 and CBP as the most probable interactors of GRA28 (Fig. 4C). Additionally, IP experiments in which HEK 293T cells expressing p300-HA were infected with either GRA28-Ty or GRA16-Ty tagged lines resulted in detection of p300 upon pulldown of the Ty epitope only when infected with GRA28-Ty parasites (Fig. 4D). CBP/p300 interacts with STAT complexes (56) and recruits DNA polymerase II to initiate gene transcription (57). We have recently shown that TgIST mediates its inhibitory effect through prevention of recruitment of the coactivators CBP/p300 to STAT1 (22). CBP/p300 acetylates numerous lysine residues on the four core histones (58), including H3K27Ac, a critical histone mark linked to actively transcribed genes (59, 60). IFNγ priming induces STAT1 recruitment and promotes H3K27 acetylation through HAT catalytic activity of CBP/p300 (61). Given the inhibitory effect of GRA28 on IFN-γ-induced transcription, we sought to investigate if this inhibition occurred through altering the HAT catalytic activity of CBP/p300. Ectopic co-expression of GRA28-Ty together with CBP or p300 in HEK 293T cells, followed by assessment H3K27 acetylation in nucleosomes from nuclear extracts resulted in attenuated acetylation of H3K27 by CBP or p300 (Fig. 5A,B). Furthermore, quantitative immunofluorescence analysis to assess acetylated H3K27 in HFFs infected with WT parasites showed decreased H3K27 acetylation levels compared to uninfected cells and cells infected with *Δgra28* parasites (Fig. 5C). This phenotype was rescued by complementing the *Δgra28* mutant with GRA28 (Fig. 5C). Collectively these findings suggest that GRA28 potentially blocks IFN-γ-driven transcription by inhibiting the catalytic activity of CBP and p300.

**Figure 5.**
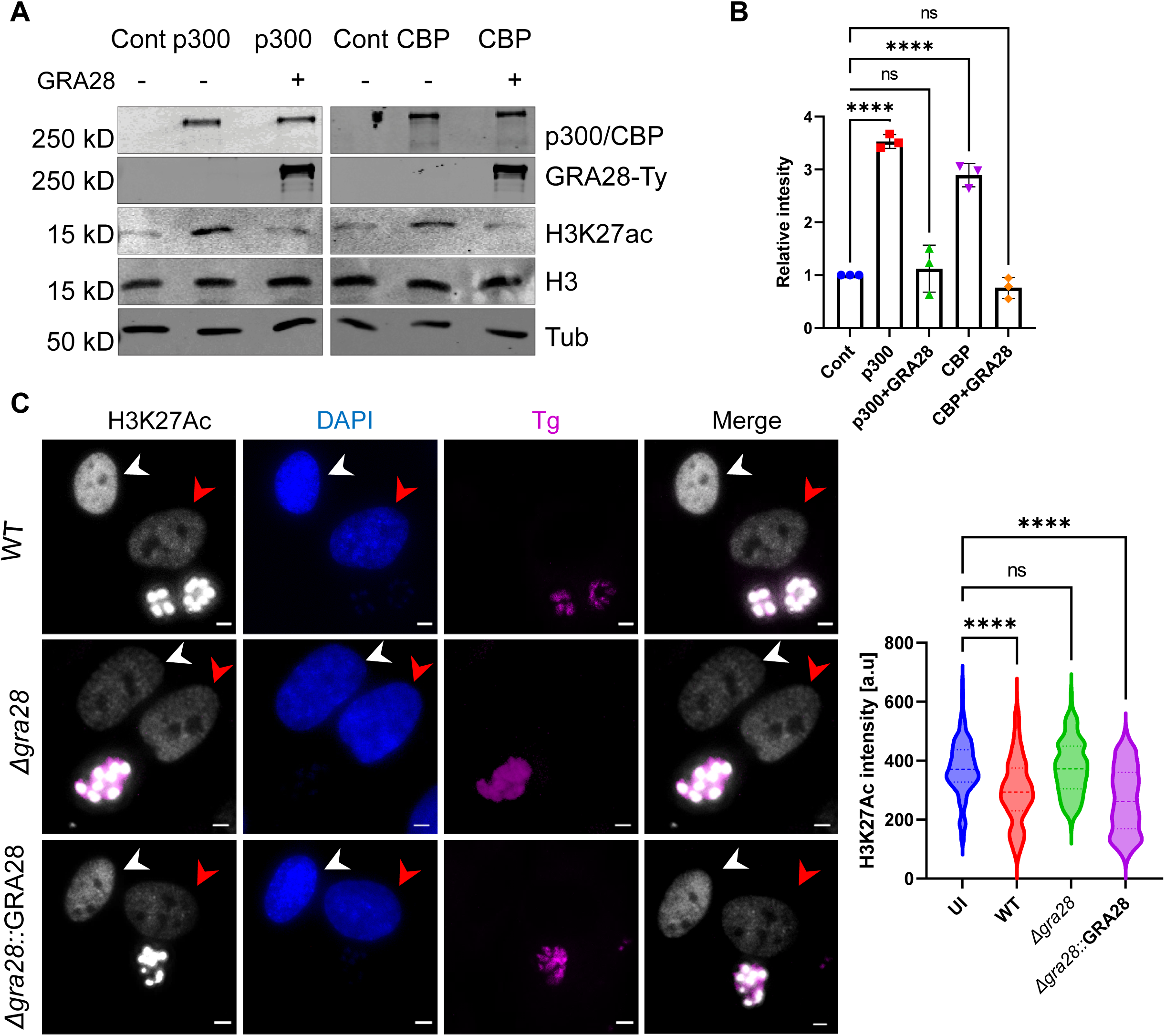
GRA28 perturbs CBP/p300 histone acyltransferase activity. (A) HEK 293T cells were transfected with CBP-HA or p300-HA in combination with GRA28-Ty or empty vector. Twenty-four hours after transfection, cells were collected lysed and fractionated into cytoplasmic and nuclear fractions. The cytosolic and nuclear extractions were resolved on SDS-PAGE gels, blotted with primary antibodies, and imaged with LI-COR specific secondary antibodies. Tubulin and Histone H3 were used as loading controls. (B) Quantification and statistics of the H3K27 acetylation (H3K27ac) in (A). Intensities of the bands corresponding to H3K27ac were measured by Image Studio then relative intensity was adjusted to H3 intensity. GRA28 driven change in H3K27ac levels were quantified from three biological replicates and is presented as a fold change between GRA28 transfected and control cells. Data presented as Mean ± SD. from three independent experiments. Statistical significance was determined using one-way ANOVA with Dunnett’s multiple comparison test. ***P<0.0001. (B) Representative images showing endogenous levels of H3K27ac in HFFs infected with RH (WT), *Δgra28* mutant or *Δgra28*::GRA28 complement. Cells were fixed 24 hr post-infection, stained with rabbit anti-H3K27ac and guinea pig anti-TgSERCA and anti-rabbit IgG Alexa Fluor 488 (white) and anti-guinea pig IgG Alexa Fluor 568 (magenta) and DAPI (blue). Scale bars = 5 µm. The graphs show the mean of nuclear H3K27ac intensity in at least 150 infected (red arrow) or uninfected cells (UI) (white arrow) per sample. Data shown are representative of three independent experiments that gave similar results. ****P < 0.0001 using one-way ANOVA test with Dunnett’s multiple comparison test.

## Discussion

IFNγ triggers an array of mechanisms that effectively limit Toxoplasma by blocking growth and/or eliminating intracellular parasites (62). The majority of the differentially regulated genes responding to *Toxoplasma* infection are MYR1 dependent (63). However, in genome wide loss of function screens none of the MYR translocon complex-dependent GRA effectors emerged as significant hits, suggesting potential collaborative functionality that obscures the individual contributions of each effector (13, 36–38, 48). In this study, we uncover that during acute stages of infection *T. gondii* relies on the MYR translocon complex to prevent host cell death and premature egress in human cells stimulated with IFNγ following infection. Inhibiting parasite egress did not halt host cell death, and this distinctive phenotype was observed in various human cell lines but not in murine cells. Through comprehensive genome-wide loss-of-function screens, we have identified TgIST, GRA16, GRA24, and GRA28 as the pivotal MYR-dependent effectors responsible for blocking the IFNγ response, thereby hindering premature egress and host cell death. Our results suggest that GRA24 and GRA28 exert direct effects on IFN-γ-driven transcription. GRA24’s transcriptional influence hinges on its interaction with p38 MAPK, whereas GRA28 associates with and disrupts the HAT activity of CBP/p300, consequently obstructing IFN-γ-driven transcription. Our findings underscore the complexity of the immune response to *T. gondii*, revealing the parasite’s sophisticated strategies for a comprehensive subversion of IFNγ signaling, extending beyond interference with the JAK/STAT1 pathway to manipulate other signaling pathways.

In murine models, the core mechanisms of IFN-γ-mediated immunity against *T. gondii* are consistently preserved across various cell types, such as embryonic fibroblasts, macrophages, and astrocytes, demonstrating a remarkable conservation of immune defense mechanisms (26–28, 64–66). However, the known restriction mechanisms in humans display significant variability among different cell types, lacking a universally applicable one. This leads to a notable difference between genes critical for protecting the parasite against the IFNγ response in human and murine cells and mice (13, 36, 37, 40, 41). The phenomenon of premature egress and host cell death, observed here in various human cell types, whether primary or immortalized, appears to be unique to nonimmune human cells. Given this contrast, it raises the intriguing question of whether TgIST/GRA16/GRA24/GRA28 impact host pathways that are either inactive or absent in human immune or murine cells. Considering *T. gondii’s* broad spectrum of hosts, exploring non-human primates and other animal lineages becomes intriguing. This would broaden our understanding of the generality of the identified effects and reveal whether the observed phenotype in mice represents a common theme or a unique adaptation.

In the current study, we have found that the premature egress phenotype in HFFs, triggered by IFNγ following infection, is tightly linked with host cell death. Remarkably, Compound 1 inhibition of PKG failed to prevent the death of infected cells in this context, which stands in stark contrast to IFNγ primed HFFs, where Compound 1 treatment effectively averted host cell death (36). Our recent screening of IFNγ induced ISGs unveiled that RARRES3 induced premature egress in *T. gondii*, a phenomenon rescued by Compound 1 treatment (34). Consequently, RARRES3 is unlikely to be responsible for the observed cell death phenotype in this study. The GRA57/GRA70/GRA71 complex plays a pivotal role in blocking early egress in IFNγ primed HFFs (36, 37), and intriguingly, the components of this complex ranked among the top 200 hits in the current CRISPR screens. However, the most significant impact on parasite survival in these screens was attributed to the MYR translocon components. This observation aligns with our ability to recapitulate the *Δmyr1* mutants’ phenotype only upon the deletion of four distinct secreted effectors. Questions persist regarding the host pathways influenced by the combined activities of TgIST, GRA16, GRA24, and GRA28 and whether these pathways overlap with those targeted by the GRA57/GRA70/GRA71 complex.

*T. gondii* employs TgIST in conjunction with GRA16, GRA28, and GRA24, to suppress the IFNγ response and inhibit premature egress during acute stages of infection. Based on our loss of function screens and gene depletion experiments it appears that TgNSM is dispensable for IFNγ response modulation during the acute stage of *T. gondii* infection. The timing of protein export via the MYR translocon complex differs among various GRAs, with TgIST, GRA16, GRA24, and GRA28 being secreted early in acute infection(67). Among these, only TgIST remains within the host cell nucleus during the bradyzoite stage, while TgNSM, a late-secreted effector during acute infection, can also be detected in the nucleus of bradyzoite-infected host cells (21). This indicates that *T. gondii* utilizes two distinct sets of effectors, each tailored to impede the IFNγ response at different stages of differentiation, with TgIST as the predominant effector in both scenarios. Given, TgNSM secretion timing coupled with the broad reach of its target NCoR/SMRT in regulating diverse biological pathways, suggests a potential role beyond just IFNγ signaling.

The combined impact of TgIST, GRA28, GRA16, and GRA24 prompts the question of their individual roles in the complex process of modulating the host’s IFNγ response. p38 MAPK was shown to be directly involved in IFNγ signaling and in the transcriptional activation of ISGs via the GAS promoter elements (68–70). GRA24 has been shown to activate proinflammatory genes, including multiple (i.e., STAT1, IL12B, NOS2) within the IFNγ signaling pathway (53). Based on our ectopic GAS reporter expression experiments we show that GRA24 functions as an activator of IFNγ dependent GAS transcription. In addition to activating genes, IFNγ is also known to suppress the expression of multiple GAS dependent genes in host cells, which can have a negative effect on pathogen survival. Some of the genes repressed by IFNγ include those involved in nutrient acquisition, immune evasion, and intracellular survival of pathogens (71). While the combinatorial expression of all the effectors results in an overall repression in IFNγ activated GAS driven transcription it is possible that *T. gondii* also relies on GRA24 modulation of p38 function to activate certain genes that are normally repressed by IFNγ and are essential for its survival. In addition to its established role in the cellular stress response and inflammation, p38 has also been associated with the orchestration of host cell death. Recent studies have shown that pathogens can exploit p38 MAPK signaling for their advantage, either to inhibit or trigger host cell death (72). Hence, it is also possible that GRA24 functions to block IFNγ driven host cell death.

p53 is a tumor suppressor that as a response to various stressors, triggers cell death by transcribing genes involved in multiple death pathways including apoptosis, ferroptosis and necroptosis (73). Upon viral infection, IFN-α/β induces p53 expression, which in turn boosts IFN signaling through a positive feedback loop (74), subsequently triggering p53-dependent apoptosis to eliminate virally infected cells and prevent virus spread (75). GRA16 alters p53 levels through an interaction with Ubiquitin Specific Peptidase 7 (USP7/HAUSP). Considering GRA16’s limited impact on IFNγ driven transcription, its significance may lie in its influence on host cell death pathways regulated by p53 (76). Alternatively, many of the target genes that are immediately induced by IFNγ/STAT1 signaling are transcription factors such as interferon response factor 1 (IRF1), IRF9 and others that then drive the expression of secondary response genes (77) promoters of which lack functional GAS elements. Notably, some genes without GAS elements are upregulated upon IFNγ stimulation, even in the absence of STAT1 (78). The premature egress phenotype occurs within a 20-hour timeline. Consequently, it is plausible that GRA16 plays an important role in regulating the interferon secondary response and STAT1-independent genes. Comprehensive transcriptional studies involving various mutant combinations at multiple timepoints in the future will provide answers regarding the specific contributions of each effector in modulating host IFNγ signaling.

IFNγ triggers the swift and typically transient initiation of early ISG transcription through the direct binding of STAT1 to accessible regulatory elements containing GAS (61, 79). This binding prompts STATs to recruit HATs CBP/p300 resulting in the alteration of histone acetylation at nearly half of the genome-wide STAT1-binding regulatory elements (61). *T. gondii* infection inhibits histone acetylation at the promoters of primary and secondary response genes upon IFNγ activation (20, 80, 81). These effects so far have been ascribed to TgIST, that extends the duration of STAT1 occupancy on chromatin, concurrently hindering its capacity to recruit p300/CBP (22). Our findings indicate that GRA28 perturbs CBP/p300 HAT activity and blocks IFNγ transcription driven by GAS elements characteristic to IFNγ primary response genes. Our current understanding of Toxoplasma induced epigenetic changes are limited to GAS sequences in promoter regions of a select few ISGs at a single time point (19, 20, 80, 81). Changes in host chromatin modification driven by *T. gondii* infection differ between primary and secondary response genes (80, 82). It remains to be determined what the individual contribution of TgIST and GRA28 are to the global epigenetic landscape of the infected cell and their functions in primary and secondary response genes.

Collectively these findings highlight that *T. gondii* has evolved a highly complex strategy to subvert and evade IFNγ driven innate immune response. Unraveling the parasite entirety of effectors necessary for survival in IFNγ stimulated cells is critical for our understanding of the mechanisms at play during infection and this knowledge could form the basis for developing strategies to combat Toxoplasmosis more effectively.

## Materials and Methods

### Parasite and Host Cell Culture

*T. gondii* tachyzoites were serially passaged in human foreskin fibroblast (HFF) monolayers cultured in D10 medium [Dulbecco’s modified Eagle’s medium, DMEM (Invitrogen)] supplemented with 10% HyClone Cosmic calf serum (Cytiva), 10 μg/mL gentamicin (Thermo Fisher Scientific), 10 mM glutamine (Thermo Fisher Scientific). HeLa cells (ATCC CCL-2), HEK 293T cells (CRL-11268), A549 cells (ATCC CCL-185), SH-SY5Y (ATCC CRL-2266), HFF hTERT (BJ-5ta) (ATCC CRL-4001), NIH-3T3 cells (ATCC CRL-1658), MEF cells, L-929 cells (ATCC CCL-1), RAW264.7 cells (ATCC TIB-71) were maintained in D10. HCT-8 cells (ATCC CCL-244) and THP-1 cells were maintained in RPMI-1640 (Invitrogen), supplemented with 10% HyClone Cosmic calf serum (Cytiva), 10 μg/mL gentamicin (Thermo Fisher Scientific), 10 mM glutamine (Thermo Fisher Scientific). THP-1 cells were first differentiated into macrophages with 50 nM Phorbol-12-Myristate-13-Acetate (PMA) (R&D systems) for 72 hrs. PMA was washed off and cells were further incubated in their maintenance media for 48 hr before use. All strains and host cell lines were determined to be mycoplasma-negative using the e-Myco Plus Kit (Intron Biotechnology). Strains used in this study are listed in Table S4. Gene disruptants and complemented lines were generated using CRISPR/Cas9 (Shen et al., 2014 24825012), as described in the Materials and Methods details. Parasite lines generated in other studies include RHΔhxgprtΔku80 (83), RH *Δmyr1* mCherry (8), RH *Δrop5* (84) and ME49 FLuc (85).

### Plasmid Construction and Genome Editing

Plasmids were generated by site-directed mutagenesis of existing plasmids or assembled from DNA fragments by the Gibson method (86). All plasmids used in this study are listed in Table S5.

### Primers

All primers were synthesized by Integrated DNA Technologies. All CRISPR/Cas9 plasmids used in this study were derived from the single-guide RNA (sgRNA) plasmid pSAG1:CAS9-GFP, U6:sgUPRT (87) by Q5 site-directed mutagenesis (New England Biolabs) to alter the 20-nt sgRNA sequence, as described previously (88). Primers for plasmids are listed in, Table S6.

### Parasite Transfection

Following natural egress, freshly harvested parasites were transfected with plasmids, using protocols previously described (87). In brief, ∼2 × 107 extracellular parasites were resuspended in 370 μL cytomix buffer were mixed with ∼30 μL purified plasmid or amplicon DNA in a 4-mm gap BTX cuvette and electroporated using a BTX ECM 830 electroporator (Harvard Apparatus) using the following parameters: 1,700 V, 176-μs pulse length, 2 pulses, 100-ms interval between pulses. Transgenic parasites were isolated by outgrowth under selection with mycophenolic acid (25 μg/mL) and xanthine (50 μg/mL) (MPA/Xa), pyrimethamine (Pyr) (3 mM), chloramphenicol (40 mM), 5-fluorodeoxyuracil (10 μM) (Sigma), Phleomycin (Phleo) (5 µg/ml) (InvivoGen) as needed. Stable clones were isolated by limiting dilution on HFF monolayers grown in 96-well plates (Figure S3).

### Pooled genome-wide screens

The genome-wide loss-of-function screen in Toxoplasma using CRISPR/Cas9 gene editing technology was performed as described previously (48, 89) (Fig. 3B). Briefly, 500 µg of gRNA library linearized with AseI was transfected into ∼4 × 10^8^ WT or *Δgra28* Cas9-expressing parasites (47) divided between ten individual cuvettes. Parasites were allowed to infect 8 × 175 cm2 flasks of confluent HFFs, and the medium changed 24 hr post infection to contain pyrimethamine and 10 μg/ml DNaseI. Parasites were passaged upon lysis (between 48–72 h post transfection) and selection continued for four passages. After four passages in HFFs, we transferred 2 × 10^7^ parasites per tissue culture flask (4 T175 flasks total) for four passages in either naive or HFFs stimulated with 100 units of human IFNγ four hours after infection. At the last passage parasite pellets with ∼1 × 10^8^ parasites were collected for DNA isolation using DNeasy blood and tissue kit (Qiagen).

### Next generation sequencing and CRISPR screen data analysis

The libraries for next generation sequencing were generated using a 2-step PCR amplification of integrated sgRNAs. Briefly, 1000ng of genomic DNA isolated from different samples was used as template for PCR1 using forward 5’ TCGTCGGCAGCGTCAGATGTGTATAAGAGACAGCCTTCTGGTAAATGGGGATGTC AAGTT 3’ and reverse 5’ GTCTCGTGGGCTCGGAGATGTGTATAAGAGACAGGAATGACACACAGGAACTACG CG 3’ primers with PCR cycling conditions of 10s 98 °C, 20s 56 °C, 15s 72 °C for 18 cycles. PCR2 was performed using 10ng of PCR1 as template using forward 5’ AATGATACGGCGACCACCGAGATCTACAC-10bp barcode-TCGTCGGCAGCGTCAGATGTG 3’ and reverse 5’ CAAGCAGAAGACGGCATACGAGAT-10bp barcode-GTCTCGTGGGCTCGGAGATGTGTA

3’ primers with PCR cycling conditions of 10s 98 °C, 20s 72 °C, 20s 72 °C for 16 cycles. PCR2 amplicons were uniquely dual indexed with 10 bp Illumina-compatible barcodes for each sample. PCR reactions were done using Q5 Taq in a 100 μl reaction. The samples within each screen were pooled and submitted to the Genome Technology Access Center, Washington University School of Medicine in St. Louis for next generation sequencing on an Illumina NovaSeq-6000 with at least 100x coverage of 150 bp paired end reads. Basecalls and demultiplexing were performed with Illumina’s bcl2fastq2 software. The forward reads were analyzed using MAGeCK as described previously (49). Briefly, read counts for the sgRNAs sequences were normalized and counted using the count function followed by comparison between the IFNγ treated population and naive control using the test function. The data was analyzed in R using the MAGeCKFlute package (90). Modified robust ranking aggregation (RRA) algorithm was used to identify positively or negatively selected genes (Tables S1, S2). The datasets generated during this study are available at Sequence Read Archive (SRA) [SRA: SUB14094820]

### IP

For mass spectrometry samples, HFF cells were infected for 24 hr with either the parasite strain RH or RH–GRA28-Ty. Nuclear extracts were prepared using the NE-PER Nuclear and Cytoplasmic Extraction Reagent kit (ThermoFisher) on ice at 4°C. Anti-TY mAb BB2 beads bound to Protein G Dynabeads (Life Technologies) were precleared and incubated with the nuclear fractions overnight. Beads were washed and used further for MS/MS analysis. Three independent replicates were performed.

### For IP experiments followed by western blot analysis

HEK293T were transiently transfected with either p300-HA, or CBP-HA (Table S5) using polyethylenimine (PEI). Twenty-four hr after transfection, cells were infected with Toxoplasma (RH) either expressing Ty-tagged GRA16 or GRA28 for 16 hr. Cytoplasmic and nuclear extracts were prepared using the NE-PER Nuclear and Cytoplasmic Extraction Reagent kit (ThermoFisher) on ice at 4°C. Anti-TY mAb BB2 beads bound to Protein G Dynabeads (Life Technologies) were precleared and incubated with the nuclear fractions overnight. Beads were washed and Ty-tagged proteins were eluted in 100 μl IgG Elution Buffer, pH 2.0 (Thermo) for 10 min at 50°C to dissociate proteins from the antibody-conjugated beads. The elutions were immediately neutralized by addition of 15 μl neutralization buffer (1M Tris, pH 8.5). Protein samples were then prepared in Laemmli buffer containing 100 mM dithiothreitol, boiled for 5 min, separated on 7.5% polyacrylamide gels by SDS-PAGE, and transferred to nitrocellulose membranes. The membranes were blocked with 5% (wt/vol) fat-free milk in PBS and then probed with primary antibodies diluted in blocking buffer containing 0.1% Tween 20. Membranes were washed with PBS + 0.1% Tween 20, then incubated with goat IR dye-conjugated secondary antibodies (LI-COR Biosciences) in blocking buffer as indicated in the figure legends or associated method details. Membranes were washed several times before scanning on a Li-Cor Odyssey imaging system (LI-COR Biosciences). Membranes were stripped between blots by incubation in stripping buffer (Thermo Fisher) for 15 min and were then washed three times for 5 min each time with PBS.

### MS/MS analysis

For immunoprecipitation experiments Protein G Dynabeads (ThermoFisher) were placed in ammonium bicarbonate, then reduced (2 mM DTT for 1 hr at 37°C) and alkylated (10mM IAM for 20 min at 22°C in the dark). One μg of sequencing grade trypsin (Promega) was then added per sample and digestion was carried out overnight at 37°C. Of the final 120 μL digest, 60 μL was dried down and redissolved in 30 μL of 2.5% acetonitrile, 0.1% formic acid. Five μL of each digest was run by nanoLC-MS/MS using a 2 hr gradient on a 0.075mm × 250mm C18 Waters CSH column feeding into a Q-Exactive HF mass spectrometer.

All MS/MS samples were analyzed using Mascot (Matrix Science, London, UK; version 2.5.1). Mascot was set up to search SwissProt 2016_11, Homo sapiens (20,130 entries), the ToxoDB-28_TgondiiME49_Annotated Proteins 20160816 (8322 entries) and the cRAP_20150130 database (117 entries) assuming the digestion enzyme trypsin. Mascot was searched with a fragment ion mass tolerance of 0.060 Da and a parent ion tolerance of 10.0 PPM. Carbamidomethyl of cysteine was specified in Mascot as a fixed modification. Deamidated of asparagine and glutamine and oxidation of methionine were specified in Mascot as variable modifications.

Scaffold (version Scaffold_4.7.5, Proteome Software Inc., Portland, OR) was used to validate MS/MS based peptide and protein identifications. Peptide identifications were accepted if they could be established at greater than 80.0% probability by the Peptide Prophet algorithm (91) with Scaffold delta-mass correction. Protein identifications were accepted if they could be established at greater than 99.0% probability and contained at least 2 identified peptides. Protein probabilities were assigned by the Protein Prophet algorithm (92). Proteins that contained similar peptides and could not be differentiated based on MS/MS analysis alone were grouped to satisfy the principles of parsimony. Proteins sharing significant peptide evidence were grouped into clusters.

### Mass spectrometry and interactome analysis

The data sets in Scaffold (v4.7.5) were filtered with min #peptide = 2, protein threshold ≤99%, and peptide threshold ≤95%. The resulting hit lists were exported to excel and edited to match the format with the SFINX analysis tool (http://sfinx.ugent.be) (Table S3). A separate excel file containing GRA28 bait was uploaded to software SFINX website (http://sfinx.ugent.be/) together with the mass spectrometry data file.

### Luciferase Reporter Assays

HeLa cells were transiently transfected, as described above. 5x-GAS conjugated luciferase reporter construct (Table S5) was used in combination with different combinations of TgIST-Ty, GRA16-Ty, GRA24-Ty, GRA24 ΔR1/R2-Ty, GRA28-Ty expression vectors or empty vector as a control. The Renilla reporter plasmid pRL-TK (Promega) was co-transfected as a control for transfection efficiency. Twenty-four hours later, transfected cells were treated with human IFN-γ at (100 U/mL for 24 hr) and firefly luciferase activity was determined using the dual luciferase reporter assay system (Promega) on SpectraMax iD3 (Molecular Devices) multimode plate imager according to the manufacturer’s protocol. Experiments were repeated at least three times.

### LDH assays

LDH assays were performed with the CytoTox 96® Non-Radioactive Cytotoxicity Assay (Promega) according to the manufacturer’s protocol. Briefly, host cells split in 96-well plates were infected for 4 hrs with 100×10^^4^ of different parasite strains and subsequently stimulated with IFN or left unstimulated. After 22 hrs, 50 µl of cell supernatant was mixed with 50 µl of assay buffer and substrate for 30 min at room temperature. The reaction was stopped with 50 µl stop solution and absorbance was measured at 490 nm. For cell death inhibitors testing the infected cells were preincubated for 30 min with relevant inhibitor or with vehicle alone (DMSO) prior to IFN-γ stimulation.

### Live imaging experiments

Parasites were inoculated into 96-well optical bottom plates (Corning) of HFFs maintained in FluoroBrite DMEM supplemented with 10% Cosmic serum, 4 mM glutamine and 10 µg/ml gentamicin. After 4 hr of infection, the 96-well plates were washed and ±IFNγ (100 u/ml) and ±5 μM compound 1 media was added. Cells were imaged starting 5 hr post infection every 10 min for 57 hr at 37°C under 5% CO2. Live imaging was conducted with a Zeiss Observer Z1 inverted microscope (Zeiss) using a Colibri 7 LED light source (Zeiss), ORCA-ER digital camera (Hamamatsu Photonics), Plan-Neofluar 10x(NA 0.3) objective (Zeiss), and ZEN Blue image acquisition software (v2.5).

### HAT assay

HAT assay was performed based on the previously published method (93).

Briefly, HEK-293T cells were transiently transfected with plasmid constructs expressing either p300-HA or CBP-HA in combination with either GRA28-Ty expression or empty vector as a control using polyethylenimine (PEI). After 48 h, cells were harvested and lysed by the freeze-and-thaw method in 50 mM Tris⋅HCl (pH 7.5), 150 mM NaCl, and 0.1% Triton X-100 supplemented with 1 mM PMSF and one Complete Protease Inhibitor tablet (EDTA-free; Roche), and then centrifuged at 13,000 × *g* for 10 min. The supernatant was used to check the expression level of CBP and P300 by Western blot using anti-P300 and anti-CBP antibodies (Cell Signaling). Anti–β-tubulin antibody (Hybridoma bank # AB_2315513) was used for the loading control. The pellet containing the nucleus was further lysed by sonication in RIPA buffer (50 mM Tris⋅HCl, pH 7.5, 150 mM NaCl, 0.5% sodium deoxycholate, and 1% Nonidet P-40) followed by centrifugation at 13,000 × *g* for 10 min. The supernatant was assayed by western blotting for acetylated histone H3K27 using anti-H3K27Ac antibody (Cell Signaling), whereas anti-H3 antibody (Cell Signaling) was used for the loading control.

For H3K27ac detection by immunofluorescence, HFF monolayers grown on glass coverslips in 24 well plates were infected with 2.5 × 10^4^ parasites per well in D10 medium fro 24 hr. Samples were fixed in 4% formaldehyde for 10 min at room temperature. Samples were blocked for 30 min with PBS containing 10% NGS, and 0.3% Triton X-100 (VWR). Samples were incubated with 1:65 anti-H3K27ac antibody (Cell signaling) and 1:1000 anti-TgSERCA in PBS containing 10% BSA and 0.3% Triton X-100 overnight, washed four times in PBS for 5 min each, and probed with 1:1000 goat anti-rabbit Alexa Fluor 488 (Life Technologies) and 1:1000 goat anti-guinea pig Alexa Fluor 568 (Life Technologies) in PBS containing 10% BSA and 0.3% Triton X-100 for 1 hr. Samples were washed three times with PBS and nuclei were stained for 5 min with Hoechst 33342 (Life Technologies) in PBS. Samples were imaged with a Zeiss Axio Observer microscope and were analyzed in Zen lite v2.1.

### Quantification and statistical analysis

All data were collected and analyzed without blinding. Data was analyzed using Prism software (version 7.01; Graphpad). Parametric statistical tests were used when the data followed an approximately Gaussian distribution, non-parametric tests were used when populations were clearly not Gaussian. Comparisons were considered statistically significant when P values were less than 0.05. Experiment-specific statistical information is provided in the figure legends or associated method details including replicates (n), trials (N), standard error, and statistical test performed.

## Acknowledgements

We thank Sebastian Lourido for sharing the genome-wide sgRNA library; John Boothroyd for providing *Δmyr1* strains; Geoffrey L Smith for providing GAS reporter plasmid; Silvia Moreno for providing TgSERCA antibody; Drew Etheridge lab for help and reagent support; We thank the Genome Technology Access Center in the Department of Genetics at Washington University School of Medicine for assistance with genomic sequencing. The Center is partially supported by National Cancer Institute Cancer Center Support Grant P30 CA91842 to the Siteman Cancer Center and by Institute for Clinical and Translational Science/Clinical and Translational Science Award Grant UL1TR002345 from the National Center for Research Resources, a component of the National Institutes of Health, and NIH Roadmap for Medical Research. Mass spectrometry was conducted by Drs. Sophie Alvarez and Michael Naldrett, Proteomics & Metabolomics Facility (RRID:SCR_021314), Nebraska Center for Biotechnology at the University of Nebraska-Lincoln. The facility and instrumentation are supported by the Nebraska Research Initiative and by a grant from the National Institutes of Health (AI118426). This study was supported by NIH grant (AI118426) to L.D.S and UGA startup fund to A.R.

Conceptualization, A.R. and L.D.S.; methodology, A.R. and L.D.S.; investigation, B.H, A.R.; formal analysis, B.H, A.R.; writing – original draft, A.R.; writing – review & editing, L.D.S. and A.R.; funding acquisition, L.D.S. and A.R.; resources, A.R, L.D.S.; supervision, A.R. and L.D.S.

Conflicts: The authors declare no conflicts.

## Data availability

The datasets generated during this study are available at Sequence Read Archive (SRA) [SRA: SUB14094820]

---

**Figure S1.**
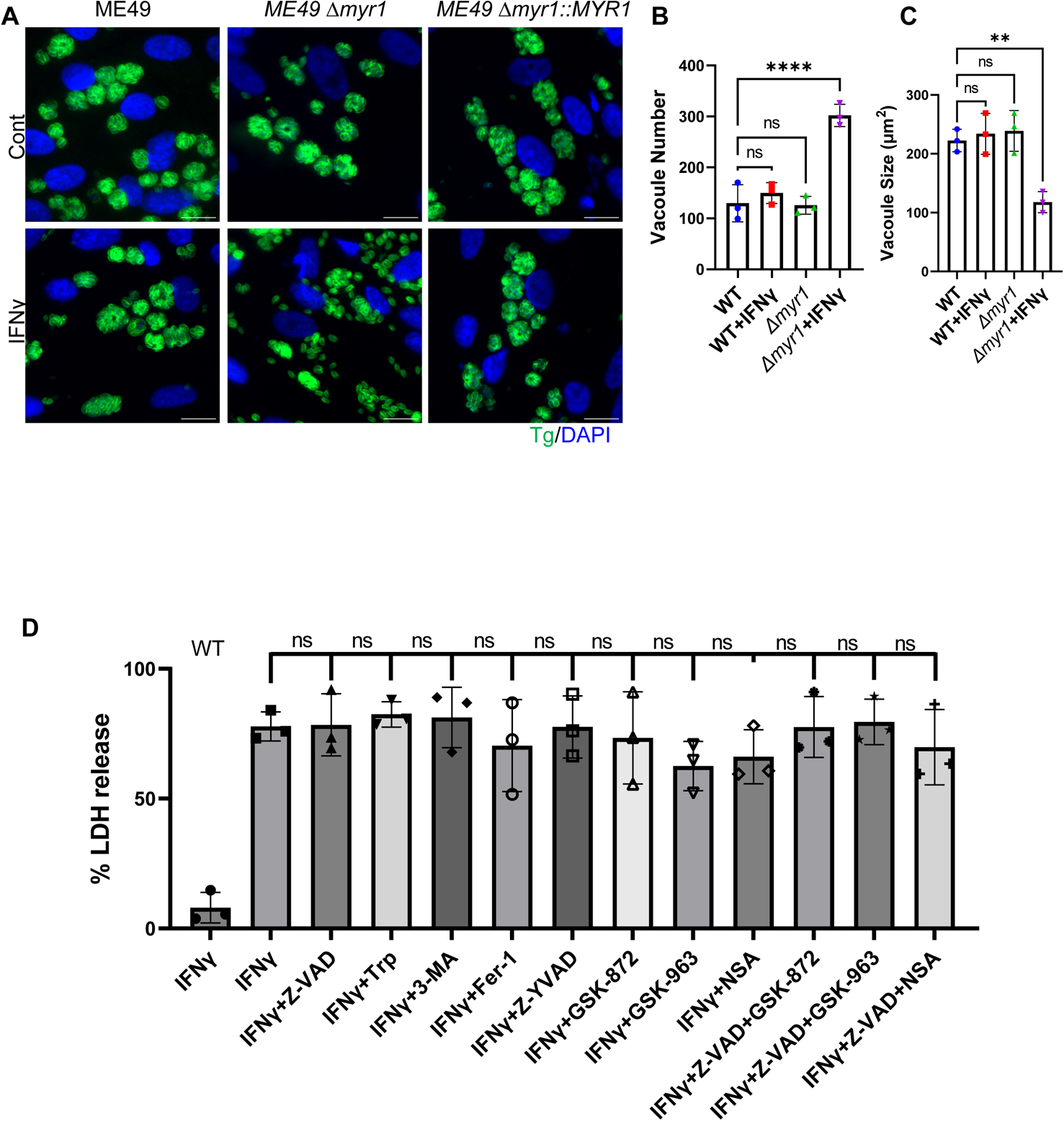
MYR1 prevents early tachyzoite egress and host cell death in type II parasites. (A) Representative images of HFF cells infected with ME49 (WT), ME49 *Δmyr1* and ME49 *Δmyr1*::MYR1 complement parasites for 24 hr prior to ± IFNγ 100 U/ml treatment. Forty-eight hours post infection cells were fixed and labelled with DAPI for nuclei (blue) and anti-GAP45 for parasites (green). Scale bar = 20 µm. (B) Average PV number per field. (C) Average PV size. Data in (B) and (C) represent Mean ± SD of three biological replicates conducted in technical duplicate with at least 30 images per sample and replicate. Statistical significance was determined using one-way ANOVA with Dunnett’s multiple comparison test. ***P<0.001, ****P<0.0001.(D) HFFs were infected with RH *Δmyr1* mutants for 4 hr before IFNγ 100 U/ml treatment in presence of cell death inhibitors Z-VAD-FMK (50 µM), GSK’963 (1 µM), GSK’872 (5 µM), NSA (10 µM), Z-YVAD-FMK (10 µM), 3-MA (5mM), Fer-1 (0.5 µM) and Tryptophan (Trp) (36 µg/ml). DMSO 0.05% only were used in control wells. Cell supernatant was collected 26 hr after infection and LDH activity was determined to measure cell lysis. Plotted is the percent of LDH release compared to maximal LDH release (after triton treatment of cells). Data from three independent experiments were pooled. Mean ± SD (n = 3 experiments, each with 3 technical replicates counted in each treatment) (A, D,F). one-way ANOVA with Dunnett’s multiple comparison test.

**Figure S2.**
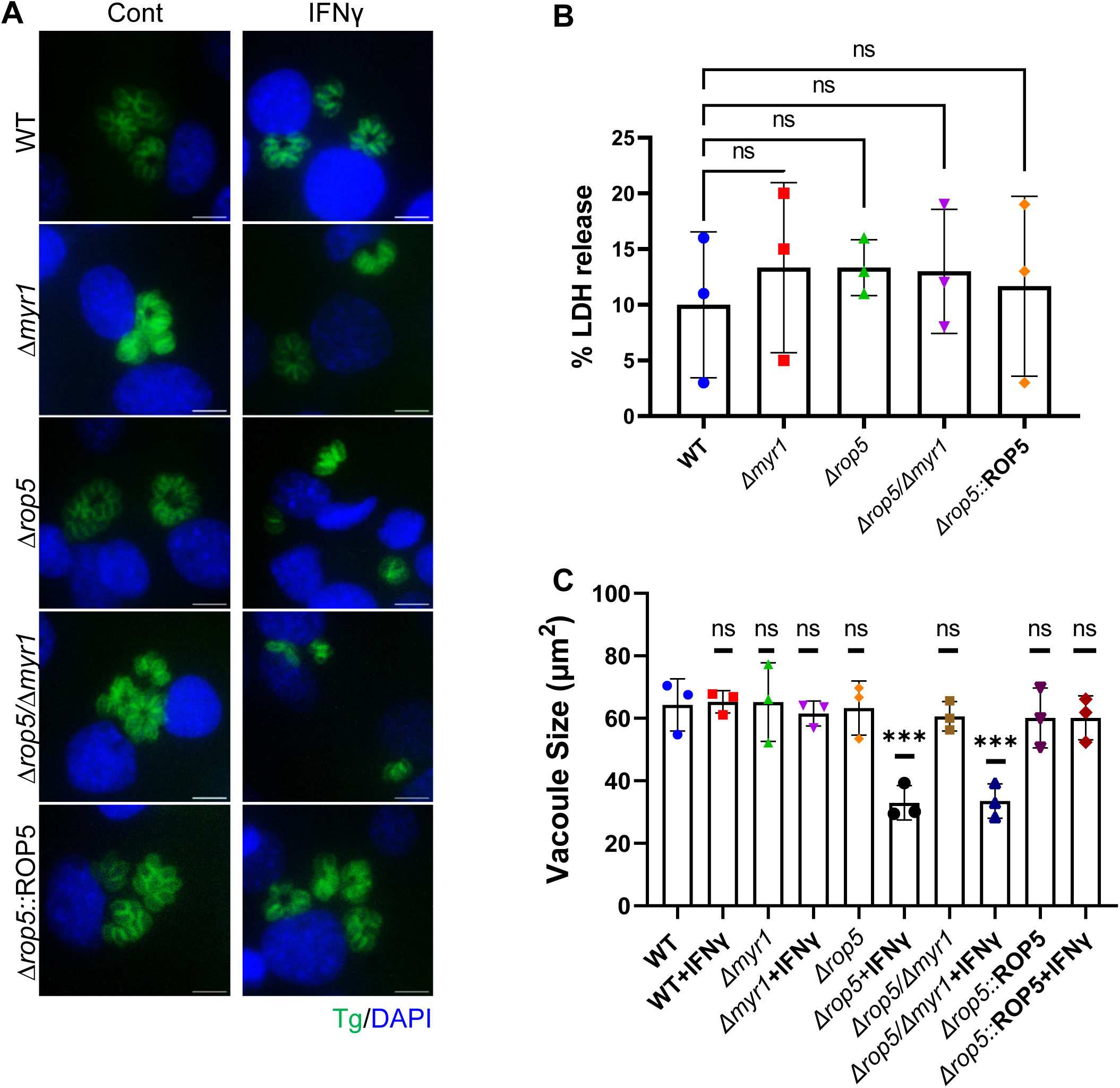
Effect of ROP5 depletion on *T. gondii* response to IFNγ murine fibroblasts. (A) Representative images of L929 cells infected with RH, *Δmyr1* and *Δmyr1*::MYR1 complement parasites for 4 hr prior to ± IFNγ 100 U/ml treatment. Twenty six hours post infection cells were fixed and labelled with DAPI for nuclei (blue) and anti-GAP45 for parasites (green). Scale bar = 20 µm. (B) LDH release in the supernatant of NIH-3T3 cells was measured 26 hr post infection. Plotted is the percent of LDH release compared to maximal LDH release (after triton treatment of cells). (C) Average PV size. Data in (B) represents Mean ± SD of three biological replicates Mean ± SD (n = 3 experiments, each with 3 technical replicates counted in each treatment). Statistical significance was determined using one-way ANOVA test with Dunnett’s multiple comparison test. Data in (C) represents Mean ± SD of three biological replicates conducted in technical duplicate with at least 30 images per sample and replicate. Statistical significance was determined using one-way ANOVA with Dunnett’s multiple comparison test. ***P<0.001.

**Figure S3.**
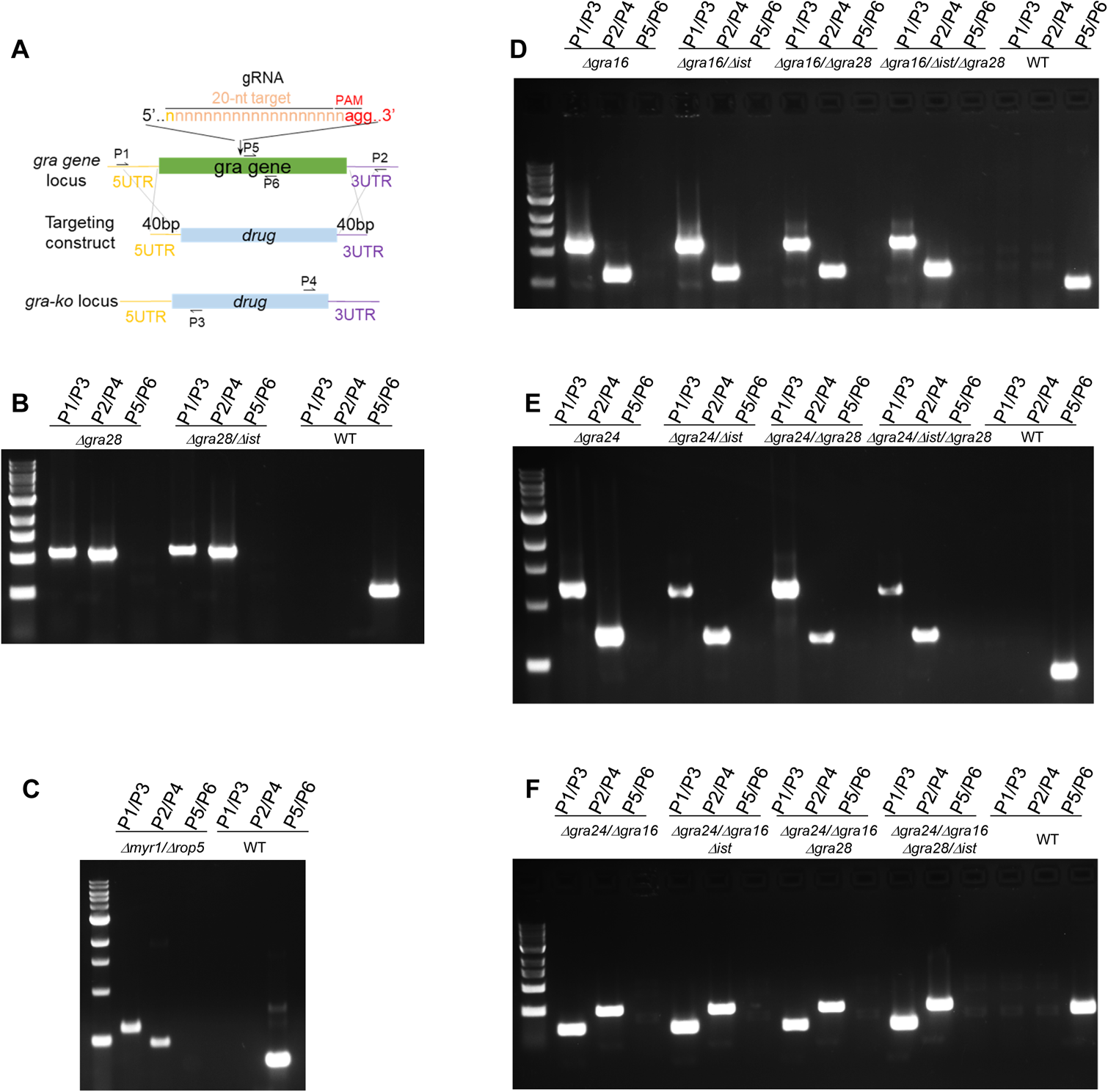
Strategy for generation of knockout lines. Schematic representation of the strategy for CRISPR/Cas9-mediated gene deletion used to generate transgenic strains used in this study. A single sgRNA expressing CRISPR/Cas9 (gRNA) plasmids targeting the middle of the genes were used to mediate double strand break and facilitate isolation of knockouts by homologous recombination. Targeting constructs consisted of a selection cassette (B) *Δgra28-*hxgprt, (C) Δmyr1-dhfr, (D) *Δgra16-*cat (E) *gra24-*cat and (F) *gra24-*phleo and short homology flanks (∼40bp) immediately upstream of the translation initiation site (left arm) and downstream of the stop codon (right arm) as homologous arms to the flanking regions of the gene of interest. Diagnostic PCRs results (B-F) to verify locus disruption. The priming sites for PCR primers are indicated: P1/P3 and P2/4 confirm integration of left and right homologous arms, respectively; P5/6 examines the integrity of the endogenous gene. A successful knockout clone gave positive PCR products with P1/P3 and P2/4 but no product with P5/6, whereas the wild type parasites gave the opposite. See also Table S6 for oligonucleotide sequences.

## Notes

### Competing Interest Statement

The authors have declared no competing interest.

